# Standardising Cellulose Production From *Acetobacter diazotrophicus*

**DOI:** 10.64898/2026.07.28.740989

**Authors:** Shalini Verma, Shiwani Kumari, Jassi Goyal, Archna Pandey, Ashutosh Sahi, Sivasakhti Ekambaram, Rimpy Kaur Chowhan

## Abstract

Bacterial cellulose (BC) is a biopolymer that comes from natural sources. It has high purity, crystallinity, mechanical strength, and biocompatibility, which makes it suitable for various biomedical and industrial uses. Unlike plant cellulose, BC does not contain lignin or hemicellulose. This results in better material quality and simpler processing. However, producing BC on a large scale is limited by high production costs, expensive culture media, and dependence on a few bacterial strains. It is essential to find cost-effective production methods and alternative microbial sources to expand its commercial use.

This study looked at the cellulose-producing ability of *Acetobacter diazotrophicus*, a safe and relatively unexplored bacterium, under different growth conditions. We compared bacterial growth and cellulose production using Hestrin–Schramm (HS) medium, the standard for BC production, and LB supplemented with glucose (LB+Glucose), which we explored as a more affordable option. We analyzed growth rates, inoculum age, and pH levels to find the best conditions for cellulose production. We observed faster bacterial growth in HS medium, with a doubling time of 3.906 hours, compared to 6.241 hours in LB+Glucose medium. Cellulose production was greatly affected by inoculum age, with successful synthesis from cultures that were agitated for 36 to 40 hours. The highest cellulose yield was at pH 6.0 in HS medium (4.6 mg/mL) and at pH 5.5 in LB+Glucose medium (3.6 mg/mL). FTIR analysis confirmed the presence of characteristic functional groups of bacterial cellulose. These results suggest that *Acetobacter diazotrophicus* is a promising and cost-effective option for producing bacterial cellulose and highlight the importance of medium composition, inoculum age, and pH for optimizing production.

## 1. Introduction

Cellulose is one of the most abundant natural biopolymers and serves as the primary structural component of plant cell walls. It is composed of glucose units linked through β-(1→4) glycosidic bonds, which provide strength and rigidity to plant tissues along with other structural polymers such as hemicellulose, pectin, and lignin. Although cellulose is mainly associated with plants, it is also synthesized by algae, protists, and several bacterial species (Marinho, 2025). Due to rapid population growth and increasing industrial demand, cellulose-based materials are widely utilized in energy production, tissue engineering, drug delivery, artificial skin, cartilage repair, bone scaffold, food additive, packaging film, water filtration, biosensor, paper industry, textile material, flexible electronics, paint additive, cosmetic material, acoustic membrane, vascular graft, ophthalmic material, antimicrobial composite (Blanco Parte et al., 2020; Fernandes et al., 2020; He et al., 2021; Kumari et al., 2023; Liu et al., 2023; Oliveira et al., 2022; Portela et al., 2019; Raut et al., 2023; Singhania et al., 2021; Troncoso & Torres, 2020).

Among the different forms of cellulose, bacterial cellulose (BC) has attracted particular interest because of its unique physicochemical and biological properties. Bacteria produce cellulose as extracellular polysaccharides by polymerizing glucose into β-(1→4)-glucan chains, which assemble into highly ordered nanofibers (Jacek et al., 2019; Revin et al., 2023). Both Gram-negative bacteria, such as *Gluconacetobacter, Acetobacter, Agrobacterium, Azotobacter, Pseudomonas, Rhizobium, Salmonella*, and *Escherichia*, and Gram-positive bacteria, such as *Sarcina*, can synthesize BC (Blanco Parte et al., 2020). Compared with plant-derived cellulose, BC is free from lignin and hemicellulose, resulting in higher purity, crystallinity, water-holding capacity, mechanical strength, and biocompatibility.

Despite its advantages, large-scale production of bacterial cellulose remains challenging because many conventional cellulose-producing strains, such as *Gluconacetobacter xylinus*, require expensive culture conditions, exhibit slow growth rates, and may undergo genetic instability during prolonged cultivation. These limitations highlight the need for identifying alternative bacterial sources that are economical, stable, and easy to cultivate for improved BC production.

As, species of *Acetobacter* are known to produce substantial amounts of exopolysaccharides and cellulose from sugars and organic acids, in the present study, we chose to work with *Acetobacter diazotrophicus* as a potential alternative to conventional cellulose-producing strains because of its low cost, easy availability, non-pathogenic nature, and suitability for patent-free academic and commercial applications. *Acetobacter diazotrophicus* is a Gram-negative, aerobic, rod-shaped, acidophilic bacterium belonging to the α-subclass of Proteobacteria and is commonly used as a biofertilizer (Dwivedi, 2020). Despite its agricultural importance, limited studies have investigated its ability to produce bacterial cellulose or attempted to standardize the conditions required for cellulose synthesis. Therefore, this study aimed to evaluate and optimize bacterial cellulose production by *Acetobacter diazotrophicus* under different growth conditions, such as, medium composition, inoculum age, and pH of culture media.

## 2. Materials and Methods

### 2.1 Materials Required

*Acetobacter diazotrophicus* bacteria are procured from N-freelancer fertilizer (Vijaya Agro Industries). Yeast extract, peptone, and Luria broth (LB) were obtained from HiMedia Laboratories. D-Glucose, citric acid anhydrous, and sodium phosphate dibasic anhydrous were from SRL. Distilled water, hydrochloric acid (HCl), and sodium hydroxide (NaOH) were used in the experiments. Standard laboratory techniques and equipment were followed for all microbiological procedures under conditions approved as being sterile by the laboratory.

### 2.2 Microorganism and Culture Maintenance

The Gram-negative, aerobic, nitrogen-fixing bacterial strain *Acetobacter diazotrophicus* was employed for its ability to grow under acidic conditions. Stock cultures were maintained on agar plates at 4 °C and cryopreserved as glycerol stocks (2:1) at −20 °C. All procedures were carried out under aseptic conditions in a laminar airflow cabinet.

### 2.3 Preparation of Culture Media

Determination of bacterial growth involves two types of media application under experimental conditions. Hestrin–Schramm (HS) medium was prepared by dissolving D-glucose (20 g/L), peptone (5 g/L), yeast extract (5 g/L), sodium phosphate dibasic anhydrous (2.7 g/L), and citric acid (1.15 g/L) in distilled water. Second, Luria broth medium was prepared and supplemented with glucose (3 gm/L). For both media preparations, the pH was adjusted to 6.0 before sterilization.

### 2.4 Inoculum Preparation

The glycerol stock was used to inoculate the sterile liquid medium, which was then incubated for 2 days. After standardising the inoculum, 1-5% (v/v) inoculum was used across all experiments.

### 2.5 Growth Kinetics

Before cellulose production, the growth kinetics of *Acetobacter diazotrophicus* were studied under controlled laboratory conditions. Hestrin–Schramm (HS) and LB+Glucose were prepared and autoclaved. For primary culture preparation, both media were inoculated with a glycerol stock of *Acetobacter diazotrophicus* and incubated in a shaking incubator at 150 rpm and 30 °C. After 48 hours of incubation, a secondary culture was prepared by inoculating 50 mL of each medium with the primary culture to obtain an initial optical density at 600nm (OD_600_) of 0.1. The cultures were further incubated under identical conditions for additional days.

Bacterial growth was monitored by measuring optical density (OD) at regular time intervals, and values were recorded at 0, 2, 4, 22, 24, 28, 46, 51, 70, 74, and 76 h using a spectrophotometer. OD values were recorded at each time point from aseptically collected culture samples immediately prior to measurement using sterile media as blanks. OD values were log-transformed to analyse bacterial growth over time. A growth curve was constructed using logOD values against time, illustrating four different growth phases, particularly the exponential growth phase. Doubling time was determined from this phase with the formula DT = 0.301 × (t□− t□)/(log OD□− log OD□).

### 2.6 Culture Conditions for Cellulose Production

Cultures were incubated at 30°C under both agitated and static conditions. After inoculation, the inoculated media were incubated in a shaker incubator at 120 rpm for 2 days. Subsequently, cultures were left undisturbed in an incubator for about 10 to 14 days at the same temperature, allowing time for a pellicle to form at the air-liquid interface of the culture.

### 2.7 Cellulose Extraction and Purification

Cellulose pellicles produced during incubation were collected using centrifugation at 6000 rpm for 6 to 10 minutes, then treated with a 2 N solution of NaOH to remove any remaining bacterial cells and residual medium components that may have persisted on the pellicle surface. Following treatment, the pellicles were rinsed multiple times with sterile distilled water until a neutral pH was achieved. The purified cellulose was dried at room temperature or in an oven.

### 2.8 Characterisation

The samples were dissolved in sterile water after purification; a small portion was then placed on a clean glass slide. The slide was air-dried and heat-fixed before being subjected to Gram staining utilizing crystal violet for primary staining, iodine for mordant, ethanol for decolorizer, and safranin as counterstain. A light microscope fitted with oil immersion was used to view (at 100X magnification) the presence/absence of any bacterial cells on the stained slides.

FTIR spectroscopy was used to identify functional groups found in bacterial cellulose after extraction, purification, and drying. FTIR was performed from 4000-400 cm^-1^ and was analyzed to demonstrate that characteristic peaks of absorption were detected from cellulose

## 3. Results

### 3.1 Growth Kinetics of *Acetobacter diazotrophicus*

The growth kinetics of *Acetobacter diazotrophicus* were analyzed in HS and LB+Glucose media by measuring optical density (OD_600_) over 76 hours. In both media, the bacteria exhibited characteristic lag, exponential, and stationary growth phases.

The lag phase of *Acetobacter diazotrophicus* was identified in the first 4 hours after inoculation into media based on no significant increase in log OD values (0.168 in HS media and 0.207 in LB+Glucose media) (Figure 1). In HS medium, the log OD values at 22 hours and 28 hours were 0.168 and 0.622, respectively (Figure 1a). Likewise, in LB+Glucose medium, log OD values at 22 hours and 28 hours were 0.207 and 0.556 (Figure 1b). The maximal bacterial growth in HS medium occurred between 46 hours and 70 hours, with log OD values of 0.740 and 0.755 (Figure 1a), and between 46 hours and 70 hours in LB+Glucose medium, with log OD values of 0.699 and 0.757 (Figure 1b). After 70 hours in both media, a slight decline in bacterial growth was observed, representing the commencement of the death phase.

**Figure 1:**
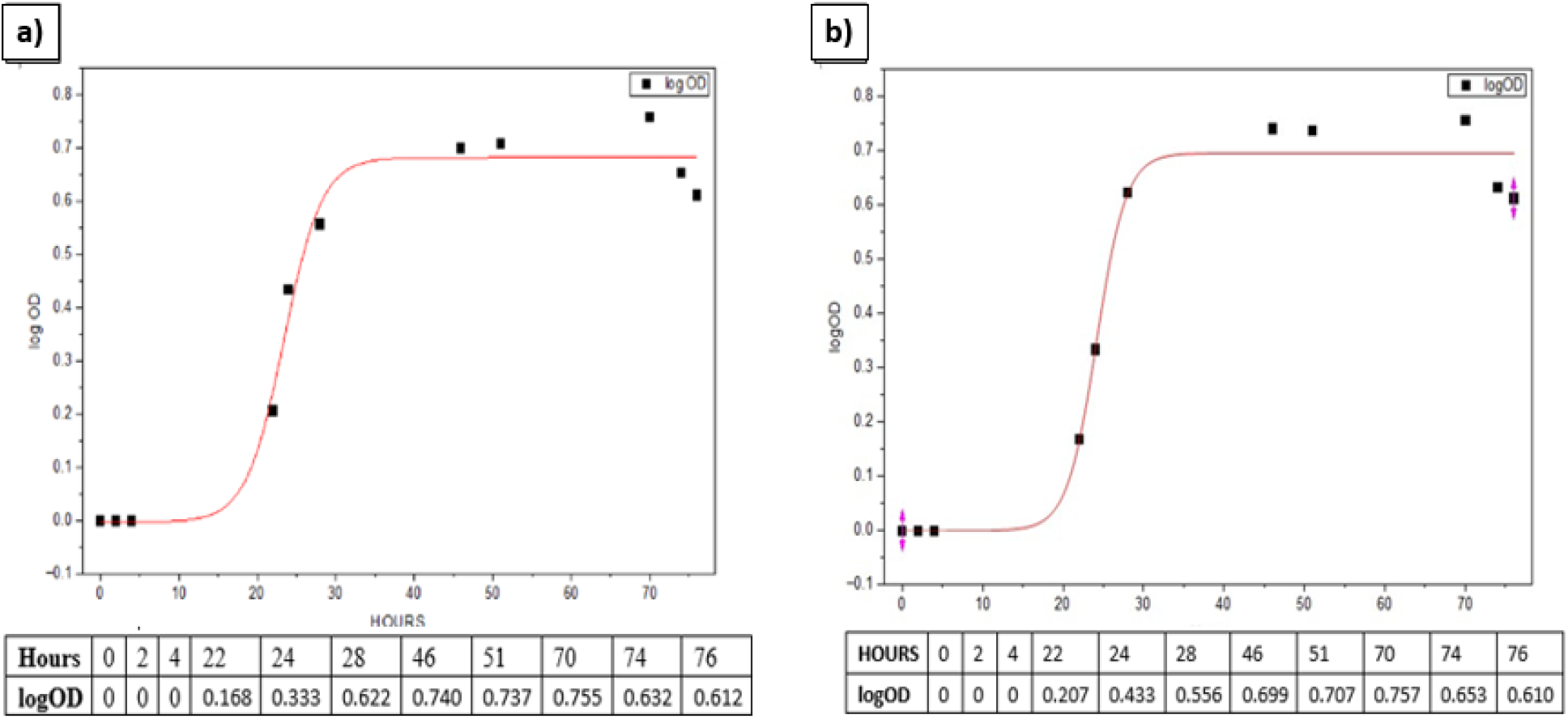
Growth kinetics of Acetobacter diazotrophicus under different culture media conditions represented by log OD values over 76 hours. (a) Growth pattern observed in HS medium and (b) growth pattern observed in LB+Glucose medium. Both media exhibited a typical bacterial growth curve consisting of lag, log, and stationary phases.

**Figure 2:**
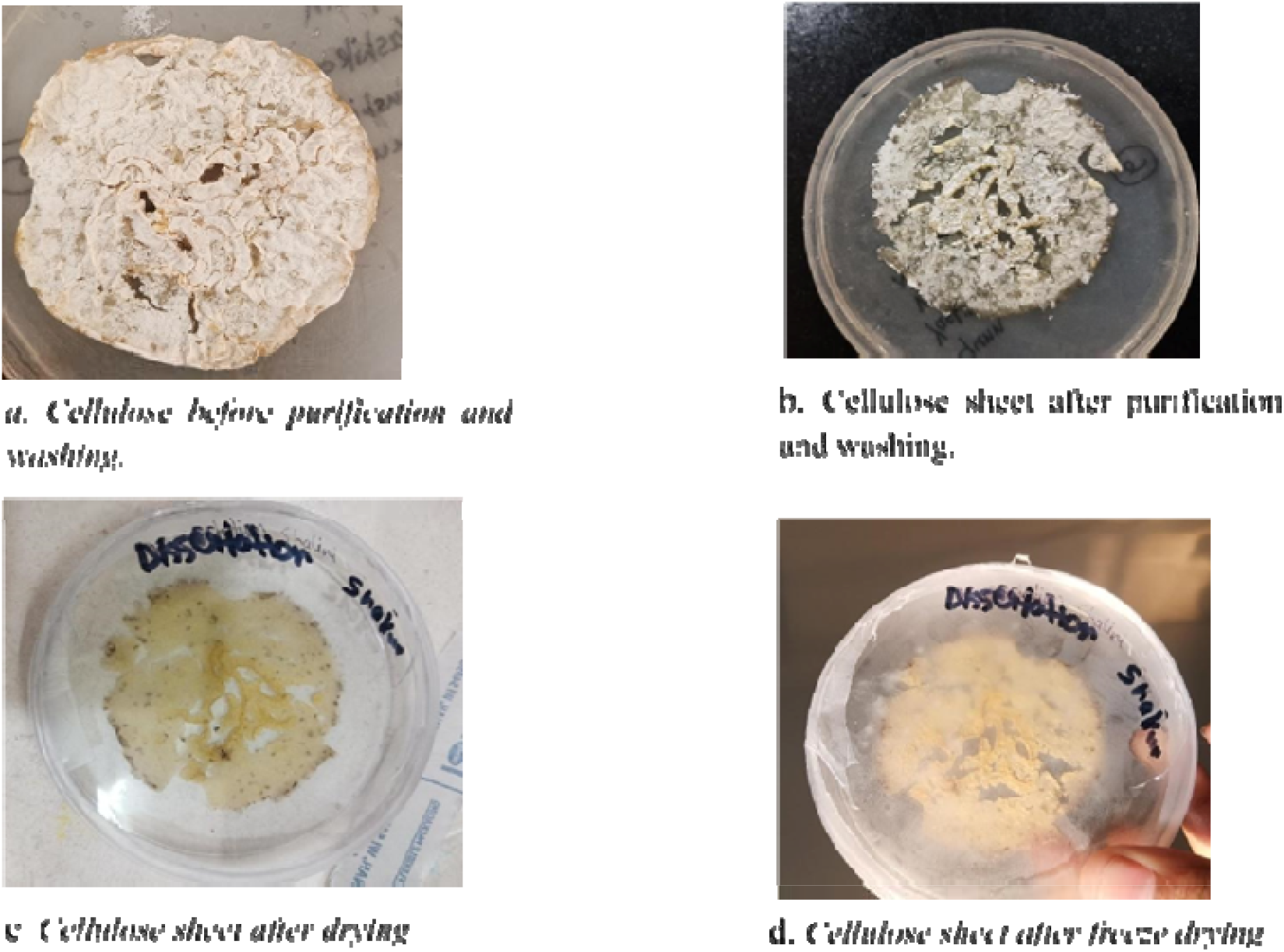
Stage of pre-cellulose processing—(a) cellulose pellicle before purification and washing, (b) cellulose sheet after purification and washing, (c) cellulose sheet after drying, and (d) cellulose sheet after freeze-drying.

The doubling times calculated during the exponential growth phase were 3.906 hours for HS medium and 6.241 hours for LB+Glucose medium. Therefore, it can be concluded that *Acetobacter diazotrophicus* grew faster in HS medium than in LB+Glucose medium.

### 3.2 Cellulose Production

#### 3.2.1 Crude Cellulose Processing

After harvesting, the recovered extract of crude cellulose intertwined with bacterial cells needs to undergo several processing steps to obtain pure cellulose. Crude cellulose that had not been purified before harvest had a yellowish tint, an irregular shape, and retained some of the bacteria and growth medium components (e.g., cells and nutrients). However, once purified with NaOH and washed with distilled water, it was visibly whiter and uniform in shape, and formed an intact sheet. Upon drying in the air, it shrank and became stiffer. When freeze-dried, it was very lightweight and porous. The dry weight of cellulose this produced, was measured to ultimately estimate the percentage yield.

#### 3.2.2 Cellulose Production at Different Inoculum Ages

The production of cellulose at different inoculum ages was evaluated in HS medium and LB+Glucose medium. For cellulose, secondary cultures were inoculated using primary cultures incubated for different agitation periods (inoculum age) to observe their effect on bacterial growth and cellulose production. Secondary culture inoculated with primary culture, agitated for 18 hours-22 hours, showed very low cellulose production. The pellet recovered after centrifugation disappeared after NaOH treatment, indicating that no cellulose was produced. In contrast, secondary cultures inoculated with primary culture agitated for 40 hours - 44 hours showed better bacterial growth and successful cellulose production.

#### 3.2.3 Cellulose Production at Different pH Levels

The production of cellulose was evaluated in HS medium and LB+Glucose medium at various pH levels (pH 5.0, 5.5, and 6.0). Differences in cellulose yield were observed among the tested pH conditions (Figure 3). The cellulose yield was calculated from purified, but not dried, cellulose pellicles. In HS medium, the maximum cellulose yield was observed at pH 6.0 (4.6 mg/mL), followed by pH 5.5 (1.6 mg/mL) and pH 5.0 (1.13 mg/mL) (Table 1). In LB+Glucose medium, the highest production was noted at pH 5.5 (3.6 mg/mL), and the lower yields were observed at pH 6.0 (1.0 mg/mL) and pH 5.0 (1.0 mg/mL) (Table 1).

**Table 1:** Cellulose yield produced by Acetobacter diazotrophicus in different culture media at varying pH conditions, incubated at 30°C. Cellulose yield was calculated as wet weight of cellulose per volume of media (mg/mL).

| Media | pH 6.0 (mg/mL) | pH 5.5 (mg/mL) | pH 5.0 (mg/mL) |
| --- | --- | --- | --- |
| HS Media | 4.6 | 1.13 | 1.6 |
| LB media | 1.0 | 3.6 | 1.0 |

**Figure 3:**
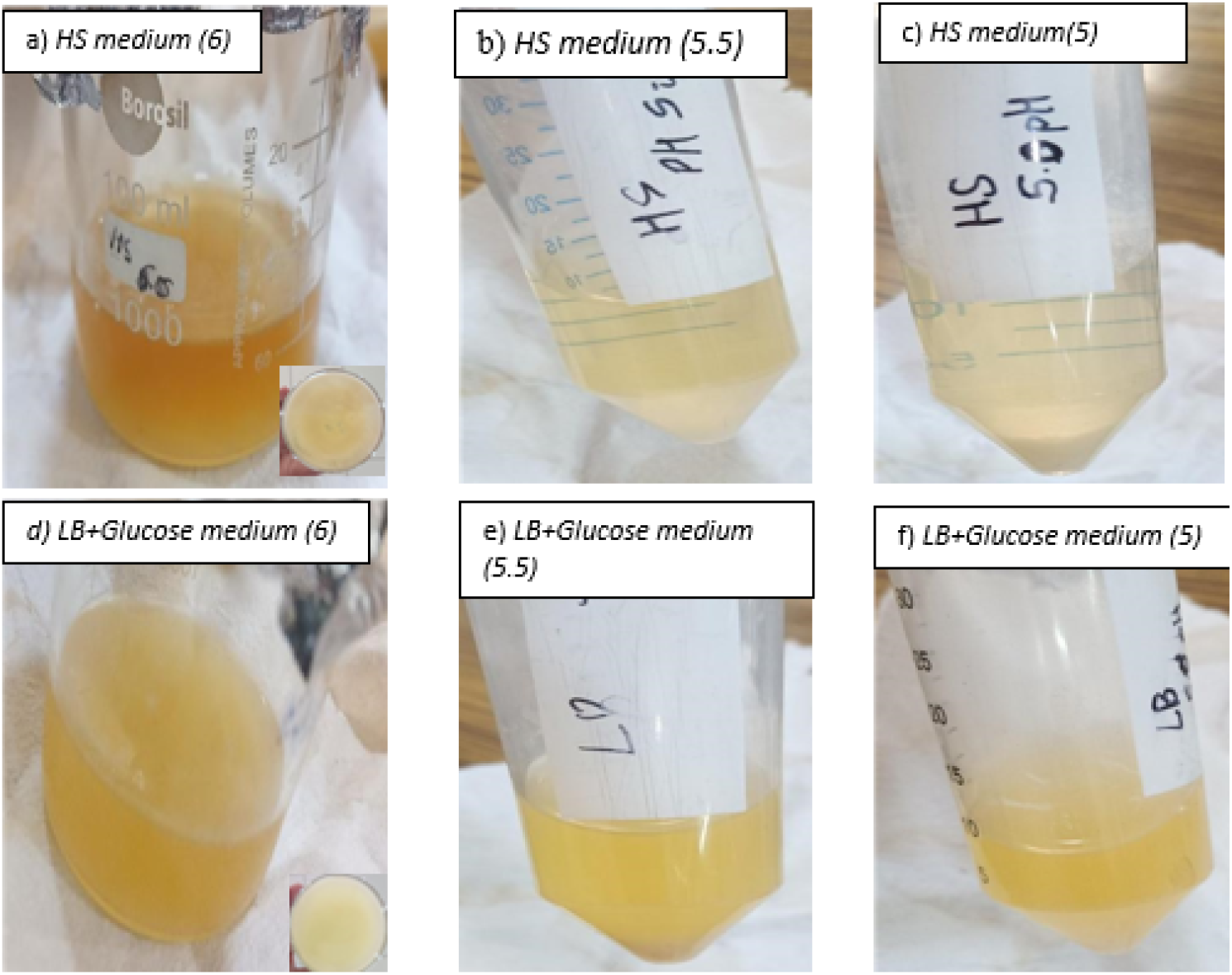
Visual observation of bacterial cellulose production in different media and pH. (a) HS medium at pH 6, (b)HS medium at pH 5.5, (c)HS medium at pH 5 (d)LB+Glucose medium at pH 6, (e) LB+Glucose medium at pH 5.5 and (f) LB+Glucose medium at pH 5.

### 3.3 Staining

A microscopic analysis of NaOH-treated samples revealed distinct thread-like cellulose structures in HS and LB+Glucose media at pH values 6, 5.5, and 5, respectively (Figure 4). The sizes of the structures were larger than the size of bacterial cells seen under the microscope, demonstrating the complete removal of bacterial biomass from the samples and the successful production of cellulose through alkaline treatment. In both media, fibres produced at pH 6 and 5.5 exhibited a greater density of cellulose than those produced at pH 5.

**Figure 4:**
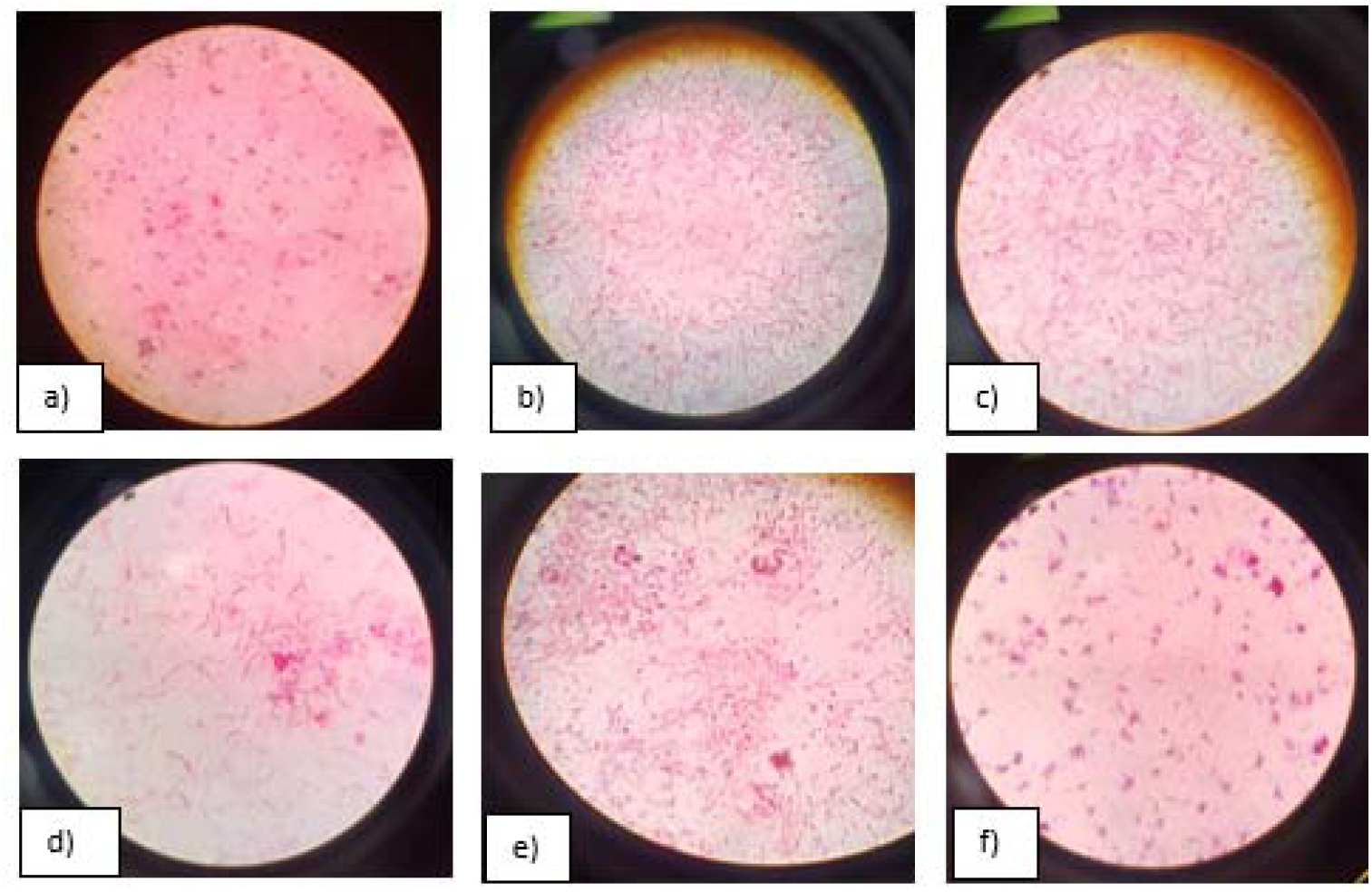
Microscopic observation of cellulose staining after NaOH treatment under different pH conditions in various culture media. (a) HS medium at pH 6, (b) HS medium at pH 5.5 (c) HS medium at pH 5, (d) LB+Glucose medium at pH 6, (e) LB+Glucose medium at pH 5.5, and (f) LB+Glucose medium at pH 5, showing the presence of purified thread-like cellulose fibres after NaOH treatment.

### 3.4 FTIR Results

FTIR analysis of cellulose samples obtained from HS medium and LB+Glucose medium was performed to confirm the presence of cellulose fibres (Figures 5 and Table 2). In both samples, broad absorption peaks were observed in the range of 3276-3291 cm□^1^ corresponding to O–H stretching vibrations, indicating the presence of hydroxyl groups. The C–H stretching vibrations were assigned to peaks around 2919 cm□^1^. The absorption band around 1636 cm□^1^ was the bending vibration of the water molecules absorbed. Other peaks identified at 1435-1413 cm□^1^ were due to CH□bending vibrations, and strong absorptions at 1058-1018 cm□^1^ were attributed to C–O–C stretching vibrations of the polysaccharide backbone. The observed FTIR spectra showed similarity with the reported characteristic peaks of bacterial cellulose, which proved that the produced samples were cellulose.

**Table 2:** FTIR peak ranges and corresponding functional groups observed in bacterial cellulose produced in HS and LB+Glucose media.

| Functional Group | FTIR Peak Range ($\text{cm}^{-1}$) | HS media | LB+Glucose media |
| --- | --- | --- | --- |
| O–H stretching vibrations | 3440–3338 <sup>[1,2,3]</sup> | present | present |
| C–H stretching vibrations | 2919–2851 <sup>[1,2,3]</sup> | present | present |
| O–H/C=O vibration | ~1662-1647 <sup>[1,2]</sup> | present | present |
| O–H and C–H bending vibrations | 1427–1361 <sup>[1,2,3]</sup> | present | present |
| C–H deformation and CH<sub>2</sub> wagging | 1335–1314 <sup>[1]</sup> | present | present |
| C–O–C stretching vibration | ~1200-1000 <sup>[2]</sup> | present | present |
| C–O and C–C stretching vibration | ~1108 <sup>[1]</sup> | present | present |
| C–O–C and C–O–H vibrations | 1054–1030 <sup>[1,2]</sup> | present | present |
| β-glycosidic linkage vibrations | 840–900 <sup>[1,2]</sup> | present | present |

Reference
<sup>[1]</sup> Akintunde et al., n.d
<sup>[2]</sup> Atykyan et al., 2020
<sup>[3]</sup> Lahiri et al., 2021

**Figure 5:**
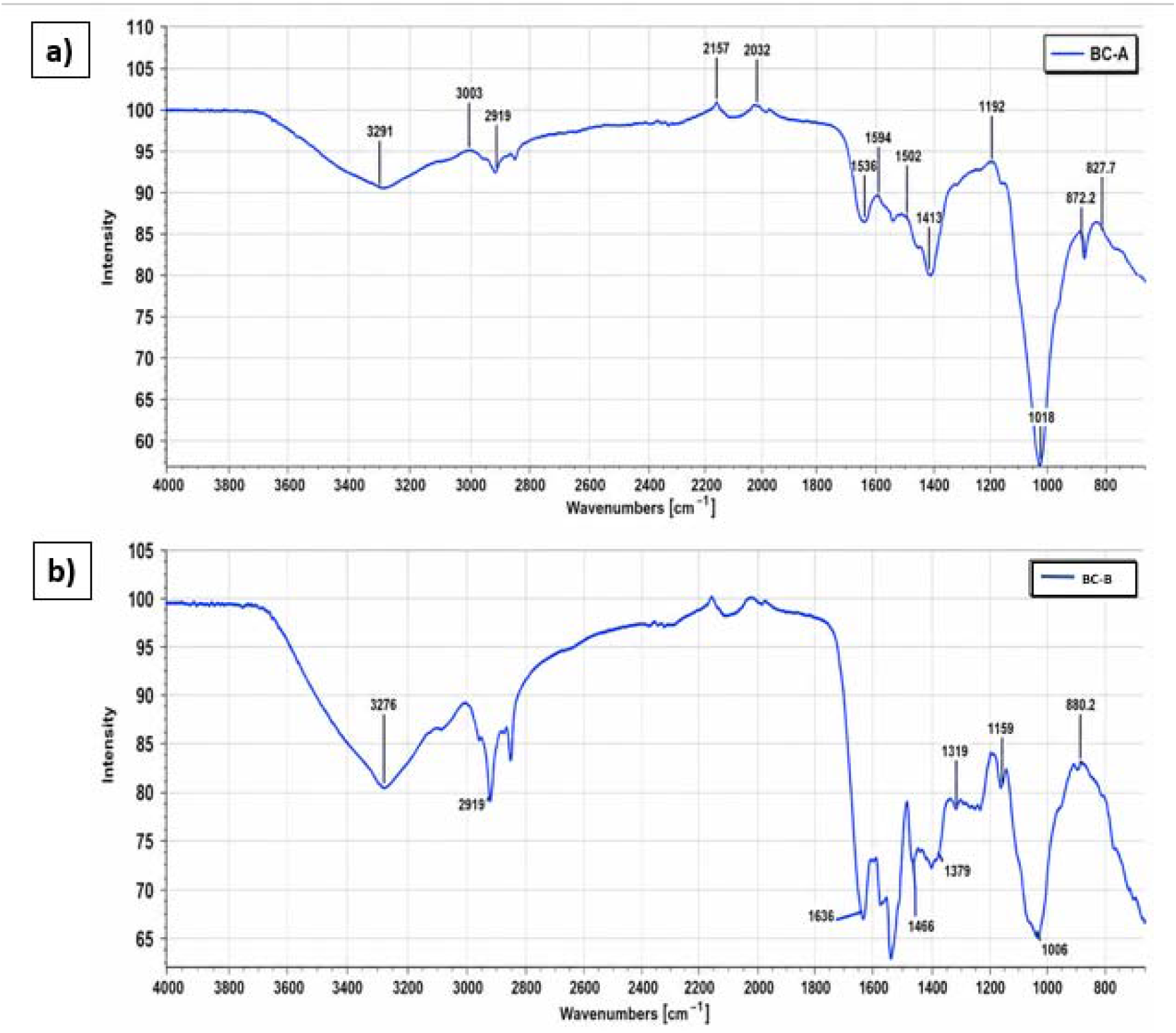
FTIR spectra demonstrating the presence of bacterial cellulose produced in different culture media. (a) Bacterial cellulose produced in HS medium (pH 6) shows characteristic functional group peaks at 3330-3400 cm□^1^ (O–H stretching), ≈2919 cm□^1^ (C–H stretching), ≈1636 cm□^1^ (H–O–H bending of absorbed water), ≈1413 cm□^1^ (CH□bending), and ≈1018 cm□^1^ (C–O–C stretching vibration). (b) Bacterial cellulose produced in LB+Glucose medium (pH 5.5) showing characteristic peaks at 3330-3400 cm□^1^ (O–H stretching), ≈2919 cm□^1^ (C–H stretching), ≈1636 cm□^1^ (H–O–H bending of absorbed water), ≈1435 cm□^1^ (CH□bending), and ≈1058 cm□^1^ (C–O–C stretching vibration), confirming the presence of cellulose in both samples.

## 4. Discussion

In the present study, the growth kinetics and cellulose production ability *of Acetobacter diazotrophicus* was evaluated under different culture conditions using HS medium and LB+Glucose medium. The results demonstrated that the composition of the growth medium, primary inoculum age, and pH significantly influenced bacterial growth and cellulose production.

The growth kinetics analysis (Figure 1) revealed that *Acetobacter diazotrophicus* exhibited a typical bacterial growth pattern consisting of lag, exponential, and stationary phases in both media. A small increase in optical density was observed during the first several hours of the incubation period, indicating a period of adaptation for bacterial cells to these new environmental conditions. The exponential phase of growth occurred between 22 hours and 51 hours in both media for *Acetobacter diazotrophicus*, with a rapid increase in optical density during this time. *Acetobacter diazotrophicus* showed faster growth in HS medium than LB+Glucose medium, as indicated by the doubling time of *Acetobacter diazotrophicus* in HS medium (3.906 hours) being shorter than that in LB+Glucose medium (6.241 hours). HS medium was designed for cellulose-producing bacteria and contains nutrients essential for bacteria to metabolise and produce cellulose. This may explain why *Acetobacter diazotrophicus* grew faster on HS medium than on LB+Glucose medium. A slight decline in growth after 70 hours suggested entry into the decline phase due to nutrient depletion and accumulation of metabolic by-products.

The findings also demonstrated that cellulose production was highly dependent on the age of the initial inoculum. Inoculations of a secondary culture using an initial culture that was agitated for 18 hours – 22 hours resulted in very little cellulose production. The pellets obtained from centrifugation disappeared after treatment with NaOH, suggesting that stable cellulose was never produced. The lack of cellulose production from early bacterial cell growth suggests that there may have been insufficient numbers of cells and that important cellulose biosynthetic pathways may not have been fully active during the exponential growth phase of each culture. In contrast, cultures inoculated using primary cultures agitated for 40 hours-44 hours showed improved bacterial growth and successful cellulose production, suggesting that inoculum taken from a culture during the later exponential phase is a better source for bacterial cellulose production.

Cellulose production was also observed to vary considerably under different pH conditions (Figure 3 and Table 1). The maximum production of cellulose was recorded at pH 6.0 in HS medium and at pH 5.5 in LB+Glucose medium. The reduced yields of cellulose at pH 5.0 in both media suggest that high acidity could have a negative impact on bacterial metabolic activity and on cellulose biosynthesis. Previous studies on bacterial cellulose-producing organisms have also reported that cellulose synthesis is strongly influenced by environmental factors such as pH, nutrient availability, and carbon source concentration(Catarino et al., 2025). The cellulase production in HS medium with a pH of 6.0 was comparatively high, and this might be the optimum condition for the cellulase biosynthetic enzymes and growth of *Acetobacter diazotrophicus*.

The cellulose nature of the obtained material in both HS media and LB+Glucose media was confirmed by staining (Figure 4) and FTIR analysis (Figure 5). The presence of fibre-like structures seen under a microscope and characteristic absorption peaks corresponding to cellulose verified the product formation. In FTIR spectra, the peaks corresponding to hydroxyl groups, C–H stretching, absorbed water molecules, and polysaccharide backbone vibrations were comparable with previously reported FTIR spectra of bacterial cellulose(Akintunde et al., n.d.; Atykyan et al., 2020; Lahiri et al., 2021) (see Table 2). The broad O–H stretching peaks observed around 3276-3291 cm□^1^ indicate extensive hydrogen bonding, which is a typical characteristic of cellulose structure. Similarly, the strong C–O–C stretching vibrations confirmed the polysaccharide composition of the produced samples. The similarity between the obtained spectra and standard bacterial cellulose spectra indicates successful production and purification of cellulose by *Acetobacter diazotrophicus* under the tested conditions.

## 5. Conclusion

The present investigation highlighted that *Acetobacter diazotrophicus* grows and produces bacterial cellulose effectively under optimized cultivation medium. Growth kinetics showed that the organism exhibited characteristic lag, exponential, and stationary phases in both HS and LB+Glucose media. The shorter doubling time observed in HS media promoted faster growth of bacterial cellulose, as compared to that in the LB+Glucose medium.

The study further revealed that, for maximum yield, inoculum age and environmental pH significantly influenced cellulose production. Secondary inoculum showed the highest yield of bacterial cellulose when agitated for 40 hours - 44 hours, compared with cultures inoculated at earlier agitation periods. These prior cultures produced negligible cellulose. Whereas, for the pH conditions, 6.0 in the HS medium and 5.5 in the LB+Glucose medium produced the most favourable outcomes. These distinct culture variables suggest that cellulose biosynthesis is strongly dependent on medium composition and pH.

Lastly, Fourier Transform Infrared (FTIR) spectroscopy supports the production of bacterial cellulose functional groups characteristic of cellulose. Overall, the results indicate that HS medium, optimal inoculum age, and appropriate pH levels positively influence bacterial cellulose production by *Acetobacter diazotrophicus*, and these findings may serve as a valuable reference for future research aimed at further enhancing yield.

## Abbreviations

BC: Bacterial Cellulose
HS: Hestrin–Schramm
LB: Luria Both

## Acknowledgments

We would like to express our sincere gratitude to **Dr Abhijit Mishra** from Shivaji College for his support and assistance in FTIR characterisation

## Statement And Declarations

### Author Contributions

S.V. and R.K.C. contributed to the study conception and design. S.V. performed conceptualisation, formal analysis, and data curation. S.V. and S.K. developed the methodology and wrote the original draft of the manuscript. S.V., S.K., J.G., R.K.C., and A.P. performed review and editing of the manuscript. R.K.C. and A.P. provided supervision and project administration. A.S. and S.E. contributed to material and design inputs. All authors read and approved the final manuscript.

